# Developmental dynamics of cross-modality in mouse visual cortex

**DOI:** 10.1101/150847

**Authors:** Ryoma Hattori, Takao K Hensch

## Abstract

Maturation of GABAergic circuits in primary visual cortex (V1) opens a critical period (CP), a developmental window of enhanced plasticity for visual functions. However, how inhibition promotes the plasticity remains unclear. Here, we investigated the developmental dynamics of auditory responses and audiovisual interactions in mouse V1. Modulation of V1 spiking activity by a transient sound was temporally dynamic with alternating enhancement and suppression phases. When paired with grating visual stimuli, sound-driven spike enhancement and suppression were weaker and stronger with preferred orientation than with non-preferred orientations, respectively, leading to impaired net orientation selectivity in V1 neurons. Strikingly, the net orientation selectivity was impervious to sound specifically during the CP due to equal total amounts of sound-driven spike enhancements and suppressions. This balance of spike modulations at the CP was achieved by the preferential maturation of sound-driven spike suppression. However, further maturation of sound-driven spike enhancement broke the balance after the CP. Spectral analysis of field potentials revealed the enhancement of a GABA-mediated sound-driven power suppression specifically at CP. Reduction of inhibition by 10-day dark-exposure or genetic deletion of GAD65 gene dampened sound-driven spike suppression in V1. Furthermore, acute suppression of either parvalbumin-expressing (PV) or somatostatin-expressing (SST) neurons suggested their early or late recruitments by sound, respectively. Our results point to the dampened net non-visual sensory influence as one of the functional roles of GABA circuit maturation during a developmental CP. The insensitivity of visual selectivity to sound during the CP may promote functional maturation of V1 as *visual* cortex.

## INTRODUCTION

Neural circuits of sensory cortex are shaped by sensory experience in postnatal life. This experience-dependent plasticity is heightened during an early developmental period, called critical period (CP). The properties and underlying mechanism of CP has been studied most extensively in primary visual cortex (V1). For example, brief visual deprivation of one eye during the CP dampens the cortical responsiveness to deprived eye stimulation (Espinosa & Stryker, 2012). In mice, the CP of V1 is between P21 and P35 with its peak sensitivity to monocular deprivation around P28 (Gordon & Stryker, 1996), and the plasticity gradually declines after the period and is almost absent around P110 (Lehmann & Löwel, 2008).

Cortical inhibition matures during CP (Lazarus & Huang, 2011), and accumulating evidence indicates that the maturation of cortical inhibitions triggers the opening of CP (Espinosa & Stryker 2012; Hensch, 2005). Mice lacking a synaptic isoform of GABA synthase (GAD65 KO) do not open their CP unless their cortical inhibition is rescued by benzodiazepine treatment (Hensch *et al.,* 1998; Fagiolini & Hensch, 2000; Iwai *et al.,* 2003; Kanold *et al.,* 2009). Similarly, cortical infusion of benzodiazepine opens a CP even before the normal onset of CP in wild-type mice (Fagiolini & Hensch, 2000). Other manipulations that indirectly affect the maturational state of cortical inhibition also shifts the timing of CP (Huang *et al*, 1999; Hanover *et al.*, 1999; Gianfranceschi *et al.*, 2003; Di Cristo *et al.*, 2007; Sugiyama *et al.*, 2008; Kobayashi & Hensch, 2015). The mechanism as to how mature cortical inhibitions enhance plasticity for visual functions has been speculative, but a recent study suggested that preferential suppression of spontaneous activities relative to visually evoked activities might enhance visual plasticity by increasing signal-to-noise (S/N) ratio of visual inputs (Toyoizumi *et al.*, 2013). However, the source of such noise that is dampened during CP has been unknown.

Although V1 predominantly receives and processes visual signals coming from thalamus, it also exhibits non-visual sensory activities such as auditory and somatosensory-related signals (Wallace *et al.*, 2004; Iurilli *et al.*, 2012; Vasconcelos *et al.*, 2011; Liang *et al.*, 2013; Romei *et al.*, 2009; Romei *et al.*, 2012; Rohe & Noppeney, 2016). Anatomical tracing studies revealed direct neural projections from auditory cortex to V1 (Charbonneau *et al.*, 2012; Lu *et al.*, 2014; Kim *et al.*, 2015), and transection between auditory cortex and V1 abolishes sound-evoked responses in V1 *in vivo* (Iurilli *et al.*, 2012). These studies suggest that multisensory interaction already occurs between primary sensory cortical areas. However, how different sensory inputs interact in primary sensory cortex remains largely unexplored. Furthermore, it is unknown how non-visual sensory activities are regulated during early postnatal life when neural circuitry undergoes extensive experience-dependent plasticity.

In this study, we investigated the impact of sound on visual responses in mouse V1 at different postnatal ages by electrophysiology *in vivo.* We found that sound-driven modulation of visual spiking activities consists of temporally dynamic oscillatory phases and depended on the orientation of concurrent grating stimuli. This sound-driven modulation of visual response impaired orientation selectivity of V1 neurons except during CP when total amounts of sound-driven spike enhancement and suppression were equalized. We further showed how local inhibitory circuits regulate the sound impacts on V1. The developmental dynamics of cross-modality and its regulation by inhibitory circuitry hint at the mechanism underlying the inhibitory control of CP plasticity.

## RESULTS

### Temporally dynamic sound-driven spike modulations and their dependence on concurrent visual orientation

Neurons in V1 preferentially respond to specific orientation of visual stimuli (Niell & Stryker, 2008). This orientation selectivity of V1 neurons allowed us to assess the sound-driven spike modulations as a function of the magnitude of concurrent visual spiking activities in each neuron. To vary the efficacy of visual inputs to individual cells during spike recordings *in vivo*, drifting gratings of different orientation were used as visual stimuli with or without concurrent white noise (WN) sound (Figure 1A). In our studies, we presented WN sound only for the first 500 ms of 3 sec visual stimulus period to examine the influence of a transient sound on visual image.

**Figure 1.**
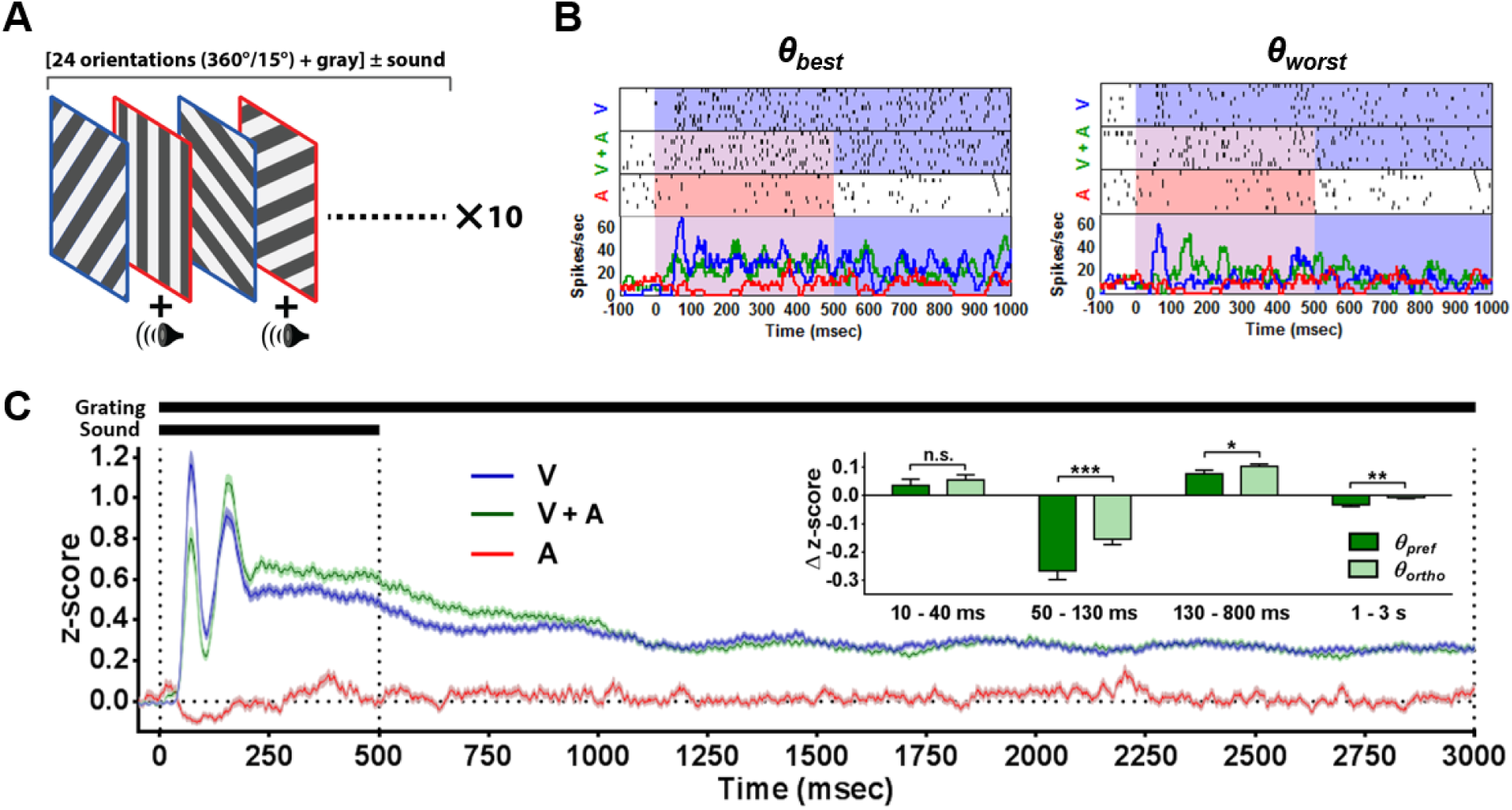
Sound-driven spike modulations and their dependence on visual stimuli in adult V1. (A) Schematic of sensory stimulation paradigm. Drifting gratings with 24 different orientations (360°/15°) were presented in randomized order alternately with or without a WN sound. ~10 trials were averaged for each condition. (B) Raster plots and peri-stimulus time histograms (PSTHs) of a typical representative cell in response to visual (V), auditory (A), and combined (V+A) stimuli. Audiovisual interactions with either the orientation that evoked the largest visual response (*θ_best_*) or with the orientation that evoked the smallest visual response (*θ_worst_*) in this example cell are shown. (C) z-scored population averages (mean ± SEM) of PSTHs (P70-90; 149 cells). PSTHs at all presented orientation were averaged for each cell. Inset shows Δ z-scores between visuo-auditory and visual conditions at each post-stimulus interval with preferred (*θ_pref_*) or orthogonal orientation (*θ_ortho_* = *θ_pref_* ± 90°) grating orientation. All error bars are SEM. Paired t-test, *P < 0.05, **P < 0.01, ***P < 0.001.

We found that the earliest sound-driven spike modulation emerges as an enhancement even before the onset of visual spiking activities (P70-90, 149 cells, peak latency of 27.2 ± 0.9 ms, mean ± SEM) (Figures 1B, 1C, 2 and S1). The fast latency of auditory response in V1 is consistent with its short latency in auditory cortex (Linden *et al.*, 2003). This transient sound-driven spike enhancement at pre-visual phase was immediately followed by a tri-phasic sound-driven modulation of visually-evoked activities, which depended on the grating orientation and persisted even beyond the sound stimulus (Figures 1B, 1C and 2). During the early (50-130 ms) and late (1-3 s) phases after stimulus onset, sound preferentially suppressed visual responses evoked by the preferred orientation. In contrast, sound preferentially enhanced visual responses evoked by the non-preferred orientation during a middle phase. The preferential multisensory enhancement with less effective orientation is consistent with the ‘inverse effectiveness rule’ of multisensory integration in classically defined multisensory areas such as superior colliculus (Stein & Wallace, 1996). However, unlike in multisensory areas, the inverse effectiveness extends to suppressive range at early (50-130 ms) and late (1-3 s) phases in V1, a classically defined unisensory area. We confirmed that the multisensory suppression is expected in areas with strong unimodality using divisive normalization model (Figure S2) which has successfully simulated multisensory integration in classically defined multisensory areas (Carandini & Heeger, 2011; Ohshiro *et al.*, 2011).

**Figure 2.**
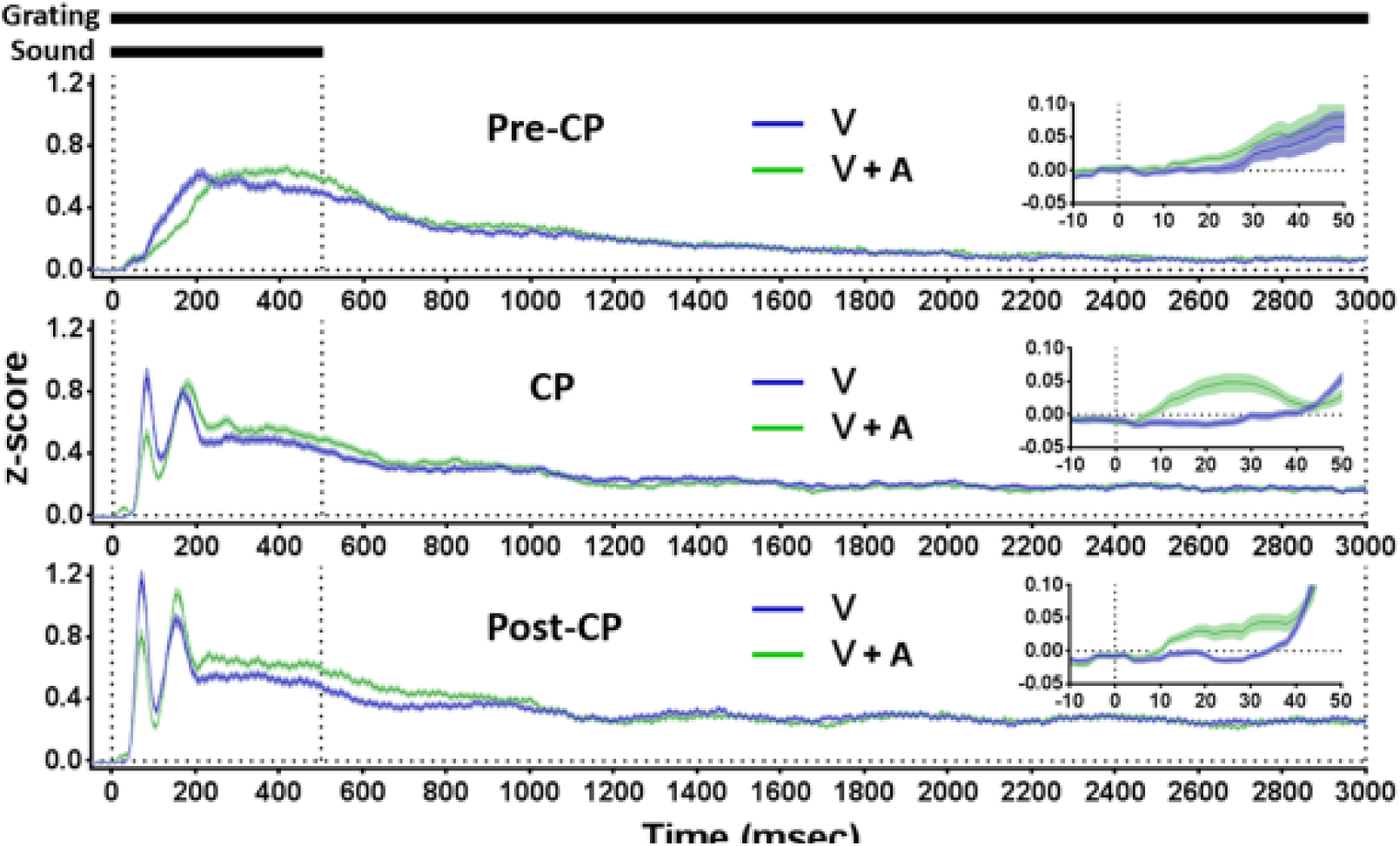
Sound-driven modulations of visual spiking activities at different postnatal periods. z-scored population averages (mean ± SEM) of PSTHs from pre-CP (P17-19, 141 cells), CP (P26-31, 108 cells), and post-CP (P70-90, 149 cells) groups. PSTHs at all presented orientation were averaged for each cell. Inset highlights the pre-visual phase.

### Sound-driven spike modulations are balanced specifically during CP

The impact of non-visual sensory inputs on V1 can be most severe when the neural circuits undergo experience-dependent plasticity for their functional maturation. Thus, we compared the sound modulations of V1 spiking activities among pre-CP (P17-19, 141 cells), CP (P26-31, 108 cells) and post-CP (P70-90, 149 cells) groups. In all age groups, a transient spike enhancement at pre-visual phase was followed by a tri-phasic modulation of visual spiking activities (Figure 2) but with different suppression/enhancement ratios (Figure 3A). The early (50-130 ms) and late (1-3 s) spike suppressions matured and reached a plateau level during CP. In addition to the magnitude maturation, the latency of early spike suppression became faster during CP (Figure 3B). On the other hand, the middle phase spike enhancement (130-1000 ms) continued to mature even after CP (Figure 3A).

**Figure 3.**
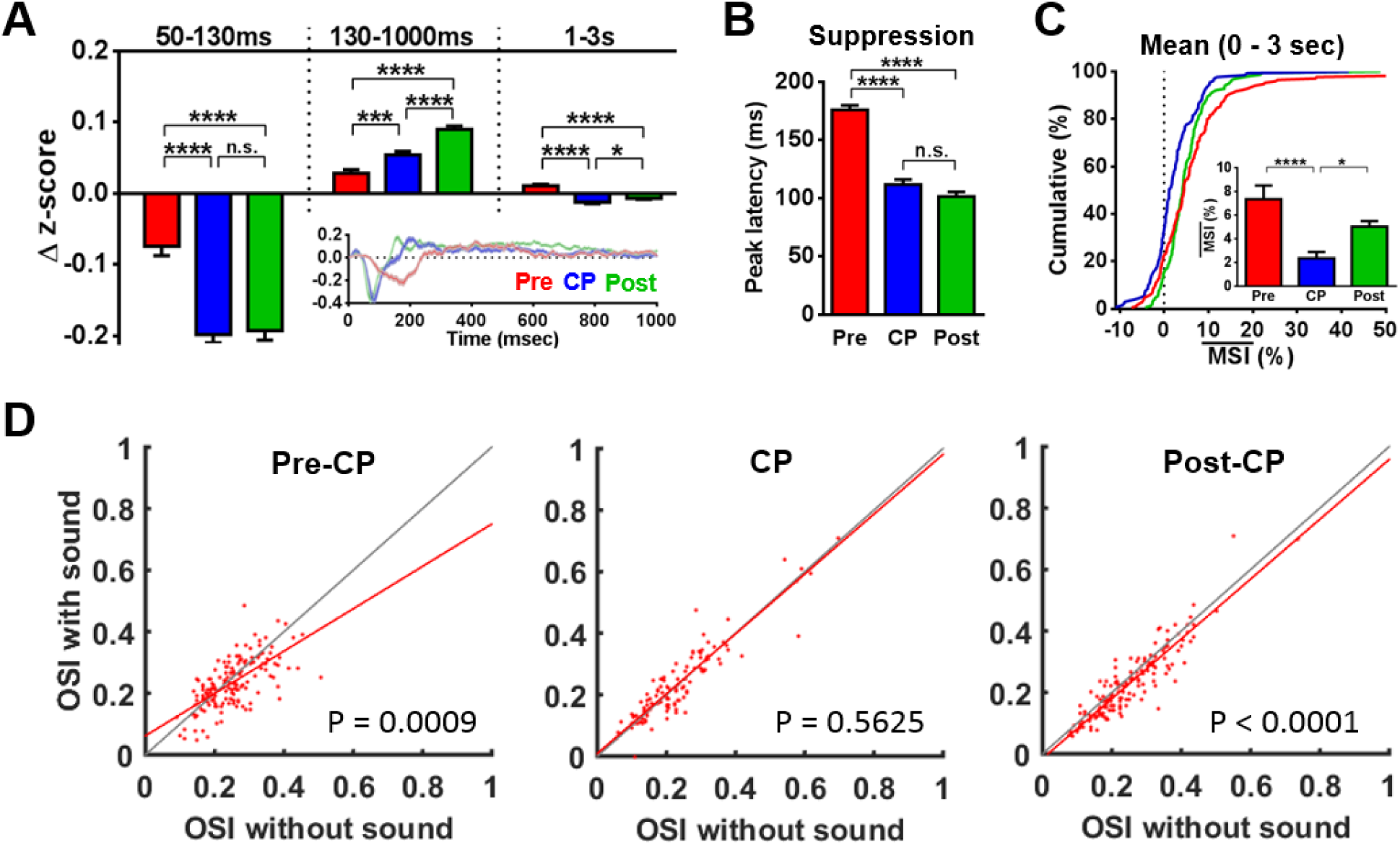
Balanced sound-driven spike modulations preserve orientation selectivity during CP. (A) Δ z-scores between visuo-auditory and visual conditions at different post-stimulus time intervals for different age groups. Inset shows the Δ z-scores as a function of time (mean ± SEM). One-way ANOVA, followed by Holm-Sidak test. (B) The peak latency of early sound-driven spike suppression. One-way ANOVA, followed by Holm-Sidak test. (C) Mean of multisensory index (MSI) at all presented orientations. Kruskal-Wallis test, followed by Dunn’s test. (D) Scatter plots indicating OSI with and without concurrent sound stimuli. Least-squares regression lines of data points are colored in red. Paired t-test. All error bars are SEM. *P < 0.05, ***P < 0.001, ****P < 0.0001.

We then examined the net impact of sound on visual response by averaging the interaction throughout the period of visual presentation (0-3 sec). We quantified the percent sound-driven modulation of visual response by multisensory index (MSI = [(VA-V)/V] × 100) (Holmes & Spence, 2005), and found that the MSI was transiently dampened during CP (Figure 3C). Sound-driven modulations of V1 spontaneous activities without concurrent visual stimuli similarly dampened their net impact during CP (Figure S1). These results indicate that sound-driven spike suppressions preferentially mature relative to enhancement during CP to offset the net modulation of V1 spiking activities.

Next, we examined the impact of sound on the orientation selectivity of V1 neurons. We found that sound impairs orientation selectivity index (OSI) of visual response in pre-CP and post-CP mice (Figure 3D). The impaired OSI is expected given the larger suppressions and smaller enhancement at preferred orientation than at non-preferred orientation (Figures 1C, S2A and S2B). On the other hand, sound did not impair OSI in CP mice (Figure 3D) due to the balanced sound-driven suppression and enhancement (Figures 3A, 3C, S1B and S1C).

### Developmental dynamics of sound-driven cortical oscillations

Having characterized the developmental dynamics of sound-driven spike modulations, we next examined the gross impact of sound on V1 neural network by recording local field potentials. We used a brief 100 ms WN sound to induce a cross-modal auditory-evoked field potential (cAEP) in L2/3 of V1. Comparison among various age groups revealed unique shifts in the average waveform morphology of the cAEP at the beginning (~P21) and after the end (~P35) of CP (7 mice for P17-18; 5 mice for P20; 7 mice for P22; 6 mice for P24; 5 mice for P27; 12 mice for P31-32; 16 mice for P71-86; 5 mice for P208-295). Before and after this CP, a characteristically large negative peak (N1) with little or no delayed positive peak (P1) was observed, while a large P1 without an early N1 emerged during the CP (Figure 4A). Spectral analysis of cAEP revealed that N1 and P1 enhances and suppresses the power of cortical oscillations at wide frequency range, respectively, and the magnitudes of power modulations correlated with the amplitudes of their peaks (Figure 4B).

**Figure 4.**
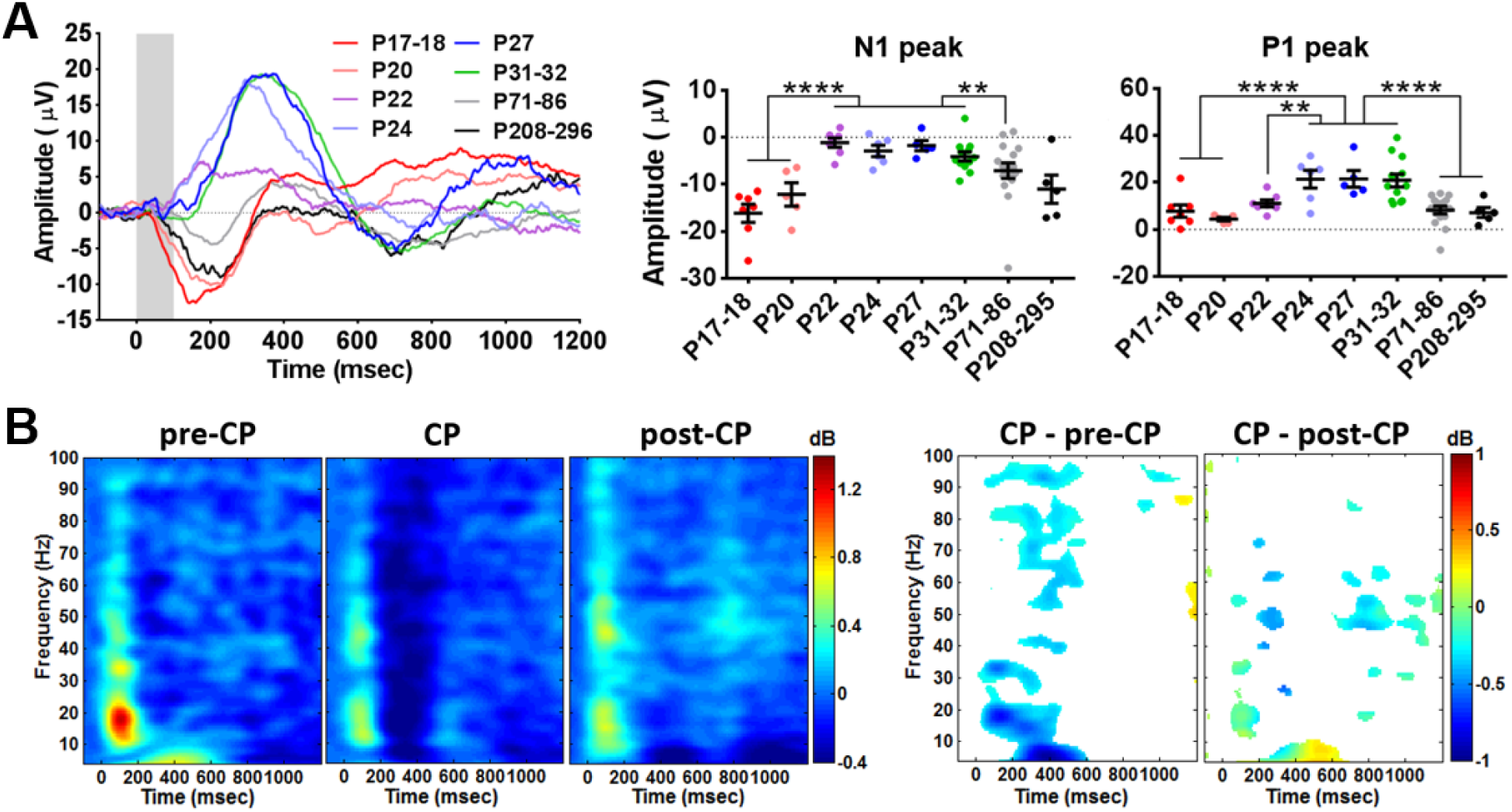
Sound-driven power suppression of cortical oscillations is specifically enhanced during CP. (A) Average cAEP traces and N1/P1 amplitudes from L2/3 across postnatal days (7 mice for P17-18; 5 mice for P20; 7 mice for P22; 6 mice for P24; 5 mice for P27; 12 mice for P31-32; 16 mice for P71-86; 5 mice for P208-295). Shading indicates WN stimulus duration. One-way ANOVA, followed by Holm-Sidak test. All error bars are SEM. **P < 0.01, ****P < 0.0001. (B) Average spectral perturbation from baseline (pre-stimulus 500ms period) after WN presentation at pre-CP (P17-20; 12 mice), CP (P24-32; 23 mice), and post-CP (> P70; 21 mice), and the statistics for the comparisons among pre-CP, CP, post-CP groups. The areas with significant difference (P < 0.05, pixel-by-pixel ANOVA, followed by Holm-Sidak test) are colored, and the color values indicate their difference in dB.

Given the maturation of cortical inhibitions in V1 during CP (Lazarus & Huang, 2011), we asked whether the sound-driven power suppression of cortical oscillations depends on GABA-mediated inhibitions. We found that the amplitude of P1 was diminished in GAD65 KO mice at CP (P27-31) (Figure 5A). However, their P1 amplitude could be fully recovered by acute local bath application of GABA-A receptor agonist, diazepam, onto V1 (Figure 5B). Further comparison of cAEPs at post-CP (P70-90) revealed the loss of P1 and enhancement of N1 in GAD65 KO mice (Figure 5C). We also recorded from mice that were exposed to complete darkness for 16-19 days from P60 (DE mice) and found the similarly diminished P1 and enhanced N1 (Figure 5C). Two weeks of dark-exposure dampens GABA-mediated inhibitions in adult rodent V1 (He *et al.*, 2010; Huang *et al.*, 2010; Stodieck *et al.*, 2014). Thus, these results suggest that inhibitory circuits regulate the relative ratio of sound-driven power suppression/enhancement in V1, and their maturation at CP is responsible for the increased sound-driven power suppression at CP.

**Figure 5.**
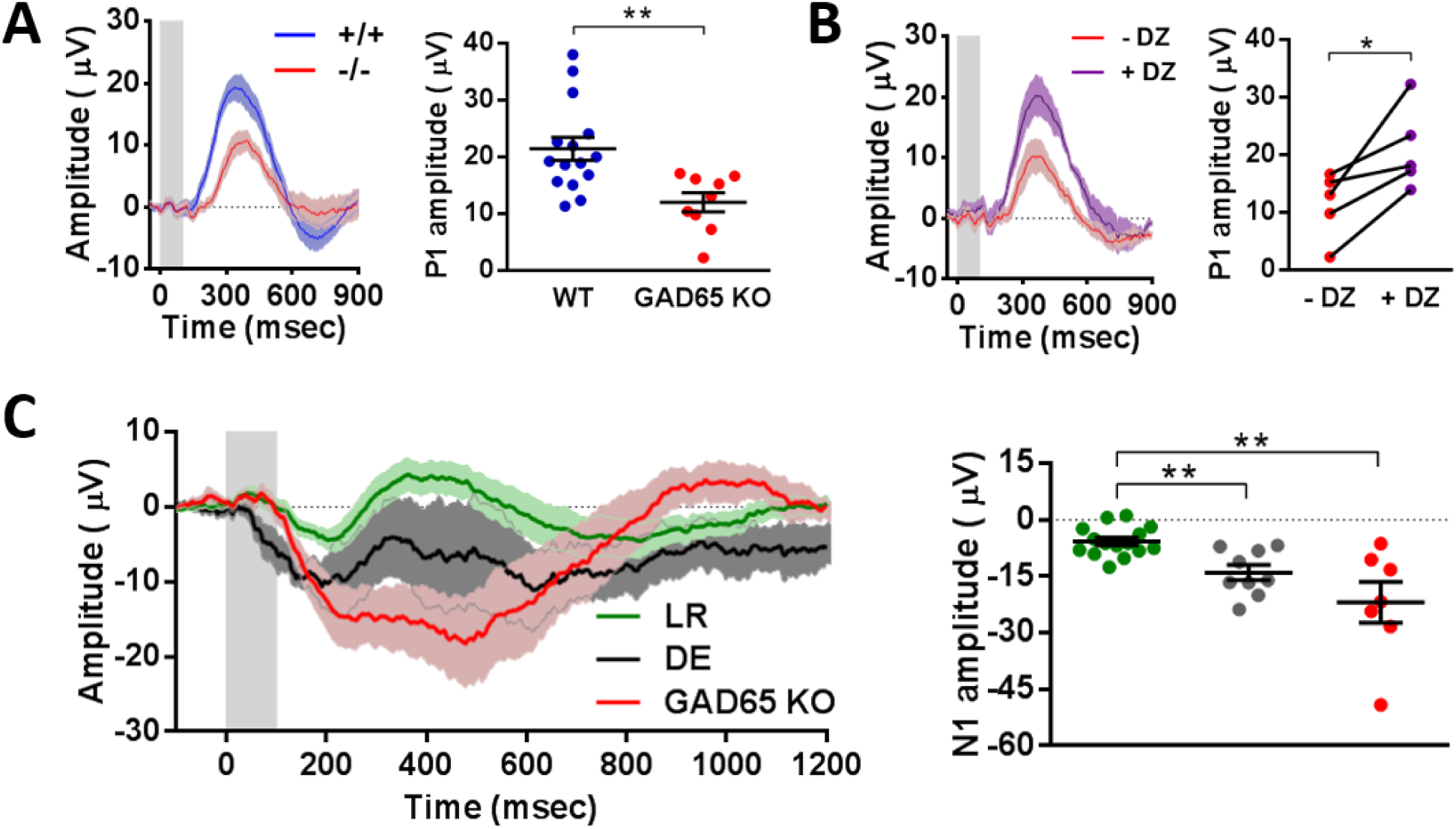
Waveform of cross-modal auditory evoked field potential depends on GABA-mediated inhibition and visual experience. (A) Average cAEP traces and P1 amplitudes from the L2/3 of WT (15 mice) and GAD65 KO (9 mice) at normal CP age (P27-31). Unpaired t-test. (B) cAEP traces and P1 amplitudes before and after acute bath application of diazepam (DZ) on the V1 of GAD65 KO mice (5 mice, P30-31). Paired t-test. (C) cAEP traces from LR (light-reared control: 15 mice), DE (dark-exposed: 9 mice) and GAD65KO (7 mice) at normal post-CP age (P70-90), and their N1 amplitudes. One-way ANOVA, followed by Holm-Sidak test. All error bars are SEM. *P < 0.05, **P < 0.01.

### Inhibitory control of sound-driven spike modulations in V1

We next examined the role of cortical inhibitions in sound-driven spike modulations in V1. First, we compared sound-driven modulations of visual spiking activities among wild-type mice with or without dark-exposure (17-18 days of dark from P60, 201 cells) and normally reared GAD65 KO mice (156 cells) at adult age (P70-90). Although the middle phase sound-driven spike enhancement (130-1000 ms) was similar across three groups, early sound-driven spike suppression (50-130 ms) was slower and weaker in both DE and GAD65 KO mice (Figures 6A and 6B). In addition, late sound-driven spike suppression (1-3 s) was significantly impaired in GAD65 KO mice (Figure 6A). These diminished sound-driven spike suppressions in DE and GAD65 KO mice increased their overall MSIs (Figure 6C).

**Figure 6.**
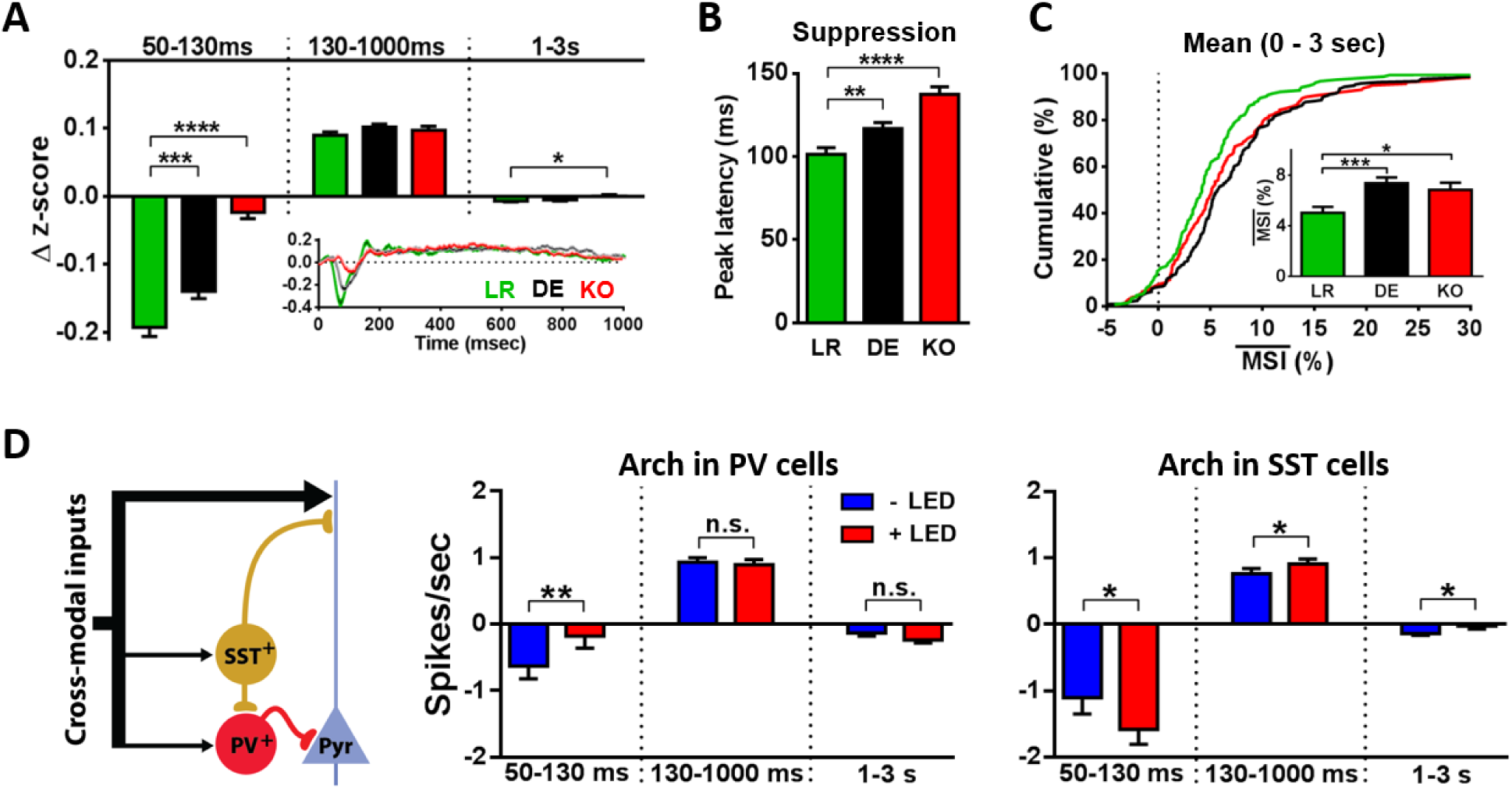
Inhibitory control of sound-driven spike modulations in V1. (A) Δ z-scores between visuo-auditory and visual conditions at different post-stimulus time intervals for LR (149 cells), DE (201 cells), GAD65 KO (156 cells) groups. All animals were recorded at normal post-CP age (P70-90). Inset shows the Δ z-scores as a function of time (mean ± SEM). One-way ANOVA, followed by Holm-Sidak test. (B) The peak latency of early sound-driven spike suppression. One-way ANOVA, followed by Holm-Sidak test. (C) Mean of multisensory index (MSI) at all presented orientations. Kruskal-Wallis test, followed by Dunn’s test. (D) Sound modulations of visual spiking activities with or without suppression of PV-expressing (140 cells) or SST-expressing (145 cells) cells. All animals (PV-Cre x Ai35, SST-Cre x Ai35) were recorded at normal post-CP age (P70-90), and suppression was performed by local LED illumination of V1. Paired t-test. All error bars are SEM. *P < 0.05, **P < 0.01, ***P < 0.001, ****P < 0.0001.

Soma-targeting parvalbumin (PV)-expressing and dendrite-targeting somatostatin (SST)-expressing interneurons are the two major inhibitory cell types and functionally mature during the CP in V1 (Lazarus & Huang, 2011). To investigate the contributions of each cell type on sound-driven modulations of visual spiking activities, we used mice expressing Archaerhodopsin (Arch), a light-gated H^+^-pump (Chow *et al.*, 2010), in either PV- (PV-Cre x Ai35) or SST-cells (SST-Cre x Ai35) to optically suppress their activities in V1 during sound presentation (Figure 6D). PV-cell suppression decreased the early sound-driven spike suppression (50-130 ms), whereas SST-cell suppression enhanced it likely by disinhibiting PV-cells (Pfeffer *et al.*, 2013). Furthermore, SST-cell suppression increased the sound-driven spike enhancement at middle phase (130-1000 ms) and decreased the late sound-driven spike suppression (1-3 s). These results suggest that the early sound-driven spike suppression is primarily mediated by PV-cells, but SST-cells mediate the late sound-driven spike suppression and limit the amount of sound-driven spike enhancement at the middle phase during audio-visual interactions in V1.

## DISCUSSION

In this study, we have characterized the audio-visual interaction and its developmental regulation in mouse V1. Through spike recordings, we have shown that sound modulates visual spiking activities by alternately repeating suppression and enhancement phases. This characteristic multisensory interaction depended on the effectiveness of visual orientation in each V1 neuron. Studies on other highly multisensory brain areas such as midbrain superior colliculus and association cortex revealed that the magnitude of multisensory enhancement is typically inversely related to the effectiveness of presented unimodal stimuli (Stein & Wallace, 1996). Similarly, we found in V1 that the sound-driven spike enhancement was smaller when paired with preferred than with non-preferred orientation for each neuron. Thus, V1 also follows ‘inverse effectiveness rule’ of multisensory integration. Furthermore, we found that the magnitude of sound-driven spike suppression was larger with preferred orientation than with non-preferred orientation. This indicates that inverse effectiveness rule can be extended to multisensory suppression. Whether audio-visual spike interactions in V1 depend on the type of sound stimuli (Romei *et al.*, 2012) and spatial disparity of cross-modal cues (Rohe & Noppeney, 2016) are subject to future studies.

We also found that the sound-driven modulations of V1 activities dynamically changes around the CP. Interestingly, net sound influence on V1 spiking activities was dampened during CP due to balanced sound-driven suppression and enhancement. As a result, the orientation selectivity of V1 neurons, which was typically impaired by sounds in pre-CP and post-CP mice, was insensitive to sounds during CP. Why does V1 dampen net auditory modulations during the CP? V1 must specialize itself as *visual* cortex by tuning and consolidating visual selectivity of each neuron during development to efficiently process visual signals later in life. Protection of visual selectivity from sound during highly plastic CP might promote this process. On the other hand, the re-emergence of net multisensory enhancement after CP might help V1 to integrate multisensory signals after establishing its identity as visual cortex. Although the weak sound-driven modulations of V1 activity reported in our study might be subconscious to animals, it can have significant impact on the maturation of visual selectivity during CP.

As a regulatory mechanism of sound-driven modulations, we identified a critical role of GABA-mediated inhibitions. Both field potential and spike recordings suggested the preferential maturation of sound-driven inhibitions during the CP. Furthermore, optical suppression of PV- or SST-cells revealed their involvement at different time course. Both PV- and SST-cells functionally mature during CP, but their maturational profiles are different. For example, PV-cells mature primarily by developing fast spiking and fast synaptic inhibition, while SST-cells increase their excitability without changing their spiking or output kinetics (Lazarus & Huang, 2011). We found that early sound-driven spike suppression, which was primarily mediated by PV-cells, becomes faster and larger during CP. Therefore, the developmental profiles of early spike suppression and PV-cell are consistent with each other.

Balanced sound-driven modulations might also contribute to CP plasticity itself. A computational model proposed that improved S/N ratio of visual modality can enhance visual plasticity in V1 (Toyoizumi *et al.*, 2013). As a potential mechanism of the improved S/N ratio during CP, the study showed that maturation of GABA-mediated inhibitions preferentially suppresses spontaneous activities relative to visually evoked activities. Our results extend the model and suggest that maturation of GABA-mediated inhibitions might increase the S/N ratio of visual inputs to enhance visual plasticity both by suppressing spontaneous activities and balancing non-visual sensory inputs. In turn, unbalanced sound-driven modulations might contribute to dampened visual plasticity after CP along with the expressions of molecular brakes for visual plasticity (McGee *et al.*, 2005; Morishita *et al.*, 2010; Pizzorusso *et al.*, 2002; Miyata *et al.*, 2012). It is worth noting that the degradation time course of visual plasticity after CP (Lehmann & Löwel, 2008) correlates well with the recovery of cAEP morphology after CP. Therefore, our findings provide a novel insight into the fundamental mechanism of circuit maturation in primary sensory cortex.

## EXPERIMENTAL PROCEDURES

### Animals

Founders of C57BL6/J mouse strain were purchased from Jackson laboratory. GAD65 KO, PV-Cre (PV-IRES-Cre), SST-Cre (Sst-IRES-Cre) and Ai35 lines were originally generated by Drs.K. Obata (Asada *et al.*, 1996), S. Arber (Hippenmeyer *et al.*, 2005), Z. J. Huang (Taniguchi *et al.*, 2011) and H. Zeng (Madisen *et al.*, 2012), respectively, and backcrossed to C57BL6/J mice. Both sexes were used. All mice were reared on 12hr light/dark cycle except during the dark-exposure of DE mice. For dark-exposure, cages were placed in a light-tight chamber within a dark room. All procedures were approved by IACUC of Harvard University or Boston Children’s Hospital.

### Field potential recordings

Mice were anesthetized with Urethane (0.8 g/kg, i.p.) / Chlorprothixene (8 mg/kg, i.m.), and then injected with Atropine (1.5 mg/kg, s.c.) and Dexamethasone (2.5 mg/kg, s.c.). A craniotomy was made over V1, and the dura was removed before placing a chamber on V1. The chamber was filled with saline. A tungsten electrode (~10 MΩ, FST) was inserted perpendicularly into V1 (~2.5 mm from lambda), and the signal was band-pass filtered (1-100 Hz). Evoked-potentials were recorded at a depth of ~250 µm for all experiments. Body temperature was maintained at 38°C, and O_2_ gas was continuously provided through a trachea tube. Both eyes were sutured shut during recordings to prevent potential visual inputs. White noise bursts (80 dB, 100 msec duration) were repeatedly presented every 2.5 sec from a speaker placed contralaterally, and ~500 trials were averaged for each animal. Acute bath application of diazepam was performed by replacing saline in a chamber with diazepam solution (14 µM in saline). We started recordings 10 min after the diazepam application.

Data analysis was carried out using MATLAB. Evoked potentials were normalized to their prestimulus period (-200 ms to 0 ms). EEGLAB toolbox (Delorme & Makeig, 2004) was used to compute sound-driven spectral perturbations from baseline. Spectral perturbations were calculated using Morlet wavelets and normalized to their pre-stimulus periods (-500 ms to 0 ms).

### Single-unit recordings

Mice were anesthetized with Urethane (0.8 g/kg, i.p.) / Chlorprothixene (8 mg/kg, i.m.), and then injected with Atropine (1.5 mg/kg, s.c.) and Dexamethasone (2.5 mg/kg, s.c.). A craniotomy was made over V1, but the dura was kept intact. Spikes were recorded (Plexon) using a linear silicon probe with 16 channels (NeuroNexus Technologies). The brain was covered by 3% agarose in saline. Body temperature was maintained at 37°C, and O_2_ gas continuously provided through a trachea tube. Drifting gratings (0.03 cycle/degree, 3 sec duration followed by 3 sec grey screen) with 24 different orientations (360°/15°) were presented in randomized order alternately with or without a white noise burst (80 dB, 500 msec duration, concurrent onset with grating) from an electrostatic speaker (Tucker-Davis Technologies) placed next to contralateral ear. Ipsilateral eye was covered by a piece of black tape to limit visual inputs to contralateral eye. ~10 trials were averaged for each condition, and the average mean rate of each stimulus condition was calculated by averaging firing rates during the period of grating presentation (3 sec). After recordings, spike waveforms were sorted by Offline Sorter (Plexon) and data analysis was performed in MATLAB.

Orientation tuning curves were drawn by fitting mean firing rates (3 sec) to double Gaussian functions with peaks 180° apart. We calculated fitting error as 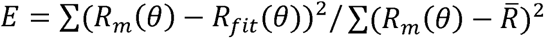 where *R_m_*(*θ*, *R_fit_*(*θ*), and 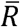, are measured firing rate at θ°, fitted firing rate at θ°, and measured firing rate averaged across all orientations, respectively (Lee *et al.*, 2012). Only units with [fitting error] < 0.6 for both with and without WN sound were analyzed. OSI was defined as *OSI* = (*R_pref_* − *R_ortho_*)/(*R_pref_* + *R_ortho_*), where *R_pref_* and *R_ortho_* are the mean firing rates of fitted tuning curves at preferred (*θ_pref_*) and orthogonal (*θ_ortho_* = *θ_pref_* + 90°) orientations, respectively.

For raster plots and PSTHs, *θ_pref_* of each cell was defined as the orientation closest to the mean of the highest peaks of Gaussian-fitted orientation tuning curves for visual and visuo-auditory conditions. MSI was calculated as *MSI* = ((*VA* − *V*)/*VA*) * 100 where *V* and *VA* are the mean firing rates (0-3 s) evoked by visual or visuo-auditory stimulus, respectively. For comparison of multisensory index among different mouse groups, we used definition of 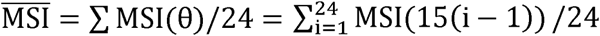 where MSIs at 24 orientations were averaged for each cell.

To draw PSTHs, firing rates were calculated using 25 ms bins with their centers 2 ms apart. Raw PSTHs were then converted to z-scores by 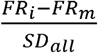 where *FR_i_* is the firing rate in the i^th^ bin of the PSTH, *FR_m_* is the mean firing rate over the entire recording period, and *SD_al_* is the standard deviation of firing rates for the entire recording period. Each z-score was further normalized to baseline by subtracting the mean baseline z-score value of each cell.

### Photo stimulation for optogenetics

An optical fiber (200 µm diameter) was positioned ~0.5 mm away from V1 and fixed by 3% agarose. Amber light (595 nm wavelength, 0.6-0.7 mW at fiber tip, Doric Lenses) was delivered through the fiber and agarose. The amber LED was turned on 0.5 s before visual stimulus onset and turned off 0.5 s after visual stimulus offset. Interleaved trials with / without amber light for each cell were compared to accurately measure cell-specific suppression.

### Simulation of divisive normalization model

We used a divisive normalization model as described previously (Ohshiro *et al.*, 2011). Weighted linear sum of visual and auditory inputs in each neuron were defined as

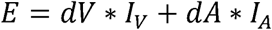

where *dV* is the dominance index for visual modality, *dA* is the dominance index for auditory modality, *I_v_* is visual input, and *I_A_* is auditory input. Unisensory inputs, (*I_v_*, *I_A_*), were given by 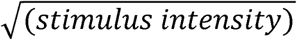 to model input nonlinearity. The activity of each neuron after divisive normalization was

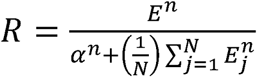

where *α* is the semi-saturation constant, *N* is the total number of neurons in the simulated network, and *n* is the exponent for the nonlinearity between membrane potential and firing rate. For our simulations, the values of *n* and *α* were fixed as *n* = 2 and *α* = 8.

Our simulation was conducted using a network of 121 cells with different combinations of *dV* and *dA* (11 x 11). *dV* in each cell takes one of 11 values from [0.5, 0.55, 0.6, 0.65, 0.7, 0.75, 0.8, 0.85, 0.9, 0.95, 1.0] while *dA* in each cell takes one of 11 values from [0, 0.01, 0.02, 0.03, 0.04, 0.05, 0.06, 0.07, 0.08, 0.09, 0.1] to simulate multisensory integration in visually dominant cortex. When V1 receives visual inputs of a specific grating orientation, some cells respond more than the others due to the heterogeneity of orientation selectivity among V1 cells. Thus, cells with large dV can be regarded as the cells in which the presented grating orientation is the preferred orientation, whereas cells with small dV can be regarded as the cells in which the presented grating orientation is non-preferred. We pooled responses of 11 cells with dV = 1 to estimate the audio-visual interaction with preferred orientation (*θ_pref_*) as the stimulus and the responses of 11 cells with dV = 0.5 to estimate the audio-visual interaction with orthogonal orientation (*θ_ortho_*) as the stimulus. Simulations were carried out in MATLAB.

## ACKNOWLEDGEMENTS

We thank M Nakamura, E Centofante and N. De Souza for mouse colony maintenance; N Picard for advice on single-unit recording. Funded by NIMH Silvio Conte Center (1P50MH094271 to T.K.H.) and the Nakajima Foundation (R.H.).

## AUTHOR CONTRIBUTIONS

R.H. designed the study, performed the experiments and simulations, interpreted the results, and wrote the manuscript. T.K.H. supervised the study.

## COMPETING FINANCIAL INTERESTS

The authors declare no competing financial interests.

## Supplemental figures

**Figure S1.**
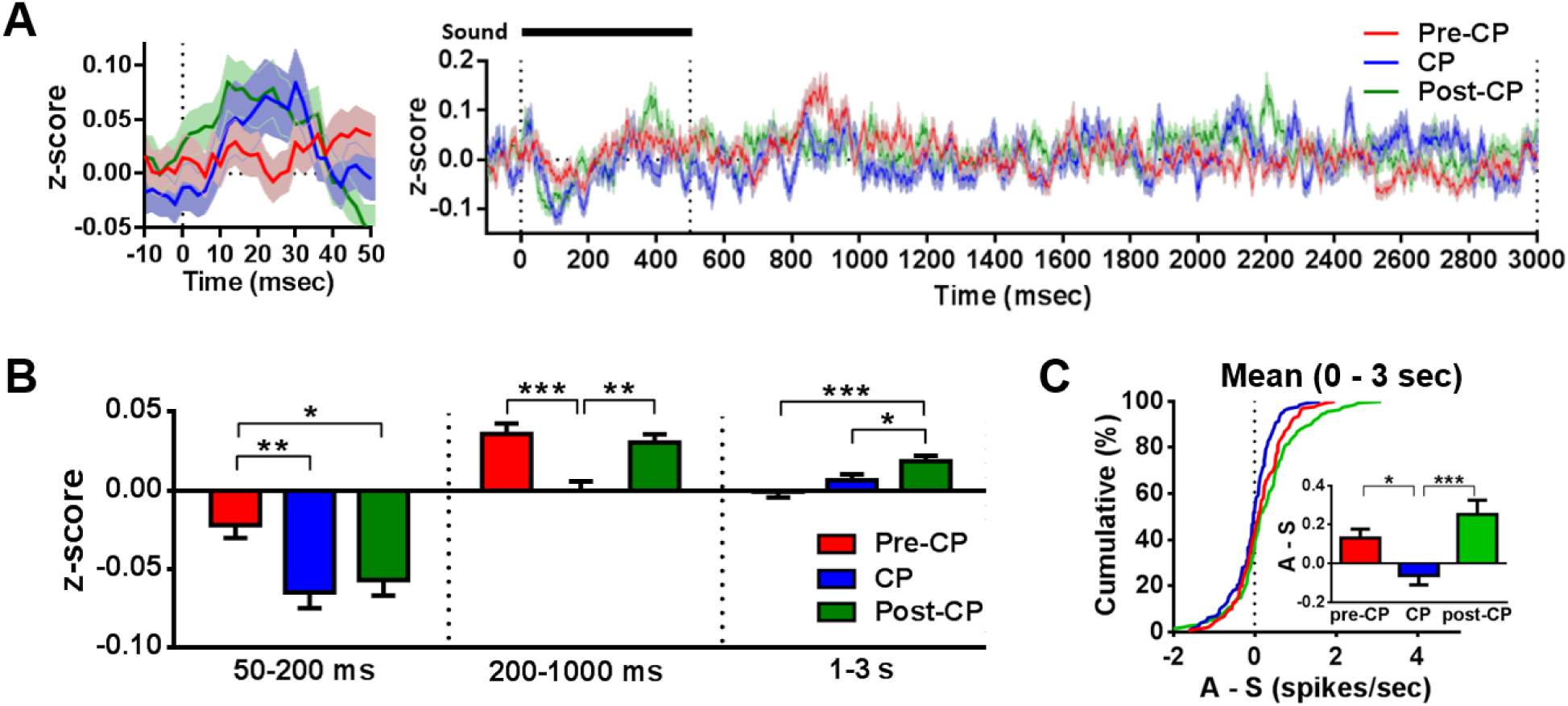
Sound-driven spike modulations of V1 without concurrent visual stimuli. (A) z-scored population averages (mean ± SEM) of PSTHs from pre-CP (P17-19, 141 cells), CP (P26-31, 108 cells), and post-CP (P70-90, 149 cells) groups. Left panel shows the magnification of -10-50 ms from the right panel to highlight the onset of sound-driven spike modulation. (B) Auditory z-scores at different post-stimulus time intervals for different age groups. (C) Sound-driven firing rate modulation relative to spontaneous activity. Average firing rate between 0-3 sec was calculated for auditory (A) and spontaneous (S) conditions, and they were subtracted from each other (A-S). All error bars are SEM. *P < 0.05, **P < 0.01, ***P < 0.001.

**Figure S2.**
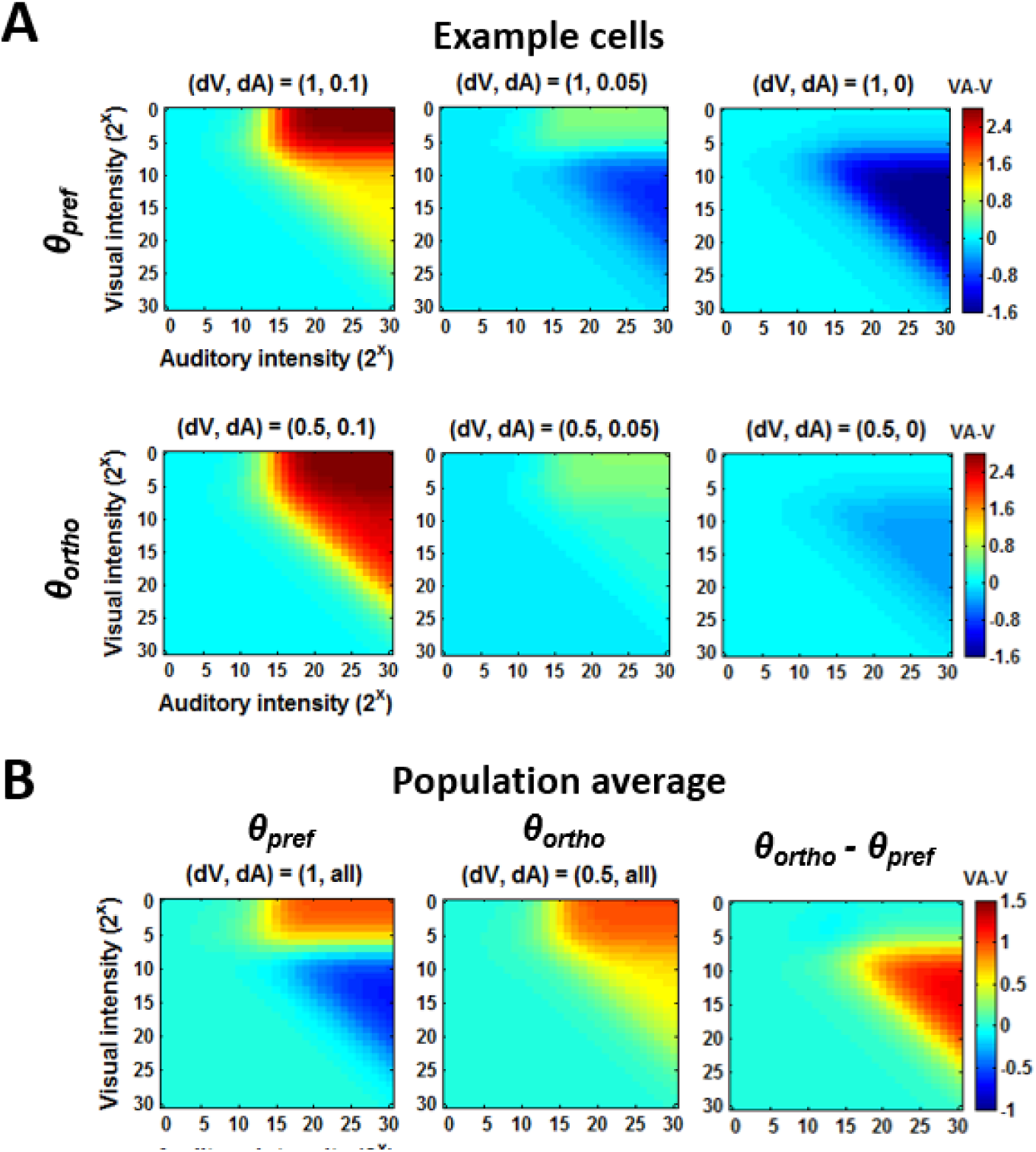
Divisive normalization model explains the dependence of audio-visual interaction on visual orientation. A network of 121 cells with different combination of visual (dV) and auditory (dA) dominance index was used to simulate audiovisual interactions in visually dominant network. (A, top row) Values of VA – V when paired with *θ_pref_* (dV = 1.0) orientation are shown for three example cells with different dA values. VA – V is shown as a function of visual and auditory stimulus intensity. (A, bottom row) Values of VA – V when paired with *θ_ortho_* (dV = 0.5) orientation are shown for three example cells with different dA values. VA – V is shown as a function of visual and auditory stimulus intensity. (B) Population average of 11 cells (dA = [0, 0.01, 0.02, 0.03, 0.04, 0.05, 0.06, 0.07, 0.08, 0.09, 0.1]) with either dV = 1 (*θ_pref_*) or 0.5. (*θ_ortho_*). VA – V with *θ_pref_* orientation and VA – V with *θ_ortho_* orientation are subtracted from each other on right panel.

